# Gut microbiota bacterial strain richness is species specific and limits therapeutic engraftment

**DOI:** 10.1101/2022.11.01.514782

**Authors:** Alice Chen-Liaw, Varun Aggarwala, Ilaria Mogno, Craig Haifer, Zhihua Li, Joseph Eggers, Drew Helmus, Amy Hart, Jan Wehkamp, Esi SN Lamousé-Smith, Robert L. Kerby, Federico E. Rey, Jean Frédéric Colombel, Michael A Kamm, Thomas J. Borody, Ari Grinspan, Sudarshan Paramsothy, Nadeem O. Kaakoush, Marla C. Dubinsky, Jeremiah J. Faith

**Author notes:** Corresponding author. (J.J.F.).

## Abstract

Despite the fundamental role of strain variation in gut microbiota function, the number of unique strains of a species that can stably colonize the human gut is still unknown. In this work, we determine the strain richness of common gut species using thousands of sequenced bacterial isolates and metagenomes. We find that strain richness varies across species, is transferable by fecal microbiota transplantation, and is low in the gut compared to other environments. Therapeutic administration of supraphysiologic numbers of strains per species only temporarily increases recipient strain richness, which subsequently converges back to the population average. These results suggest that properties of the gut ecosystem govern the number of strains of each species colonizing the gut and provide a theoretical framework for strain engraftment and replacement in fecal microbiota transplantation and defined live biotherapeutic products.

## Introduction

Strain level variation in gut microbiome composition shapes the microbiota’s influence on the host^1–6^. Understanding the strain-level structure of the microbiome could inform how best to manipulate the gut microbiome to improve health. A range of tools have enabled initial explorations of strain level gut microbiota structure, transmission, and sharing^7–14^. Despite these advances, we still lack insights into the fundamental organization of strains in the human gut microbiota including how many unique strains of a species can stably colonize the gut (i.e., the strain richness of a species).

Compared to more open ecosystems, like the ocean or soil, the gut ecosystem represents a unique semi-closed ecological niche that maintains constant temperature, diverse nutrients, and a semicontinuous unidirectional flow of nutrients. Each of these physical parameters can influence the niche availability for different strains in a species. Other factors that might influence gut microbiota strain diversity are the frequent use of antibiotics in clinical care that might reduce strain diversity in susceptible species^15^, the variation in pangenome size across species leading to accessory genomes of various sizes, host health-status, and cultural factors that might limit the consumption of sufficiently diverse sources of gut microbes to maximally fill all gut niches. Among the bacterial species commonly found in the human gut, *Bacteroides fragilis* is the only species whose strain-level structure has been well studied and it is well documented that *B. fragilis* almost always maintains a single unique strain per gut microbiota^7,16–18^.

To quantify the strain richness of common species in the gut microbiota, we performed high-throughput culture and metagenomics sequencing. With over three-thousand gut bacterial isolates from over seventy common gut species, we demonstrate that average strain richness varies across species but is low compared with other environments. In a cohort of 13 individuals receiving single-donor fecal microbiota transplantation (FMT) for the treatment of recurrent *C. difficile* infection^8^, we find that strain richness is transmissible from the donor microbiota to the recipient microbiota. In a second cohort of 56 individuals receiving FMT for the treatment of ulcerative colitis^19^, we explore the impact of supraphysiologic administration of bacterial strain richness using pooled FMT of between three and seven donor stools. In these recipients, only a fraction of the potential donor strains engrafted initially, and, by 5 years post-FMT, the average strain richness of each species had decreased to match the population average strain richness in un-transplanted individuals. These results carry immediate implications for the design of microbial therapeutics. In particular, the principle of limited species-specific strain richness suggests the potential for rationally designed microbial cocktails that consider the numbers of strains per species to use in a drug, the potential requirement for maintenance doses to maintain a high strain richness, ecological limits from resident microbial strains that may need to be removed to improve strain engraftment for some species, and the potential to use these limits to perform targeted strain replacements.

## Results

### Determining the average strain richness of gut bacterial species

To estimate the strain richness of gut bacterial species in the human gut, we obtained 3,253 cultured, sequenced gut bacteria isolated from 66 individuals (34 healthy, 23 with inflammatory bowel disease, and 9 with recurrent *C. difficile* infection; Table S1). We focused on 73 commensal gut bacterial species that were isolated from at least 3 different individuals. Pairwise genome distances from isolates of the same species (i.e., conspecific) cultured from two unrelated individuals typically have a fractional k-mer distance of at least 0.04 whereas the k-mer distances between conspecific isolates from an individual are almost always far less than 0.04 (Fig. S1A). As previously described ^20–22^, these distances are the result of most isolates of a given species in an individual being replicates of the same strain that is reisolated many times, while the same strain is very rarely shared between unrelated individuals. Therefore, we count the number of unique strains for a species (i.e., strain richness; *SR*) within an individual *i* for a given species *j* (i.e., *SR_ij_*) as the number of genomes with a k-mer distance of at least 0.04. To calculate the average strain richness for a species *j* across our study population (i.e., *SR_j_*), we average *SR_ij_* across all individuals in our cohort that harbor species *j* within their microbiota.

Across these 73 species in 66 individuals, we observed that strain richness *SR_j_* in the human gut microbiome varies significantly by species, is less than 2.0 strains per species on average (1.28 ± 0.33), and below 3.0 for all species (p=5.61 x 10^−10^, Kruskal-Wallis, Fig. 1A, Table 1). Strain richness of the commensal species *B. fragilis* was 1.0, which aligned with prior findings^7, 16, 17^. We initially sequenced, on average, 4.4 genomes per species per individual. To determine if this limited sampling depth led to a large underestimation of *SR_j_*, we sampled deeply across a subset of ten bacterial species from four major gut phyla whose estimated *SR_j_* ranged from 1.00-2.00 (Fig. S1B-D; Table S2). In addition to performing deeper culturing of the original donor sample, we also transferred some of the original human donor stools to ex-germ-free mice consuming various diets, since previous studies have demonstrated that different dietary conditions and the mouse gut environment can enrich for different strains^23–26^. After a 254% increase in sampling for these organisms, we observed 29 new strains across 509 additional isolates (20.71% increase in unique strains) suggesting we were near saturation of the strain richness for these species. The new strains belonged to species with higher *SR_j_* (Fig. S1E-F). To test the generalizability of these results, we calculated *SR_j_* for species from a previous study by Poyet et.al., where 1947 gut bacterial isolates were cultured from healthy 11 individuals^27^ (22.57 mean genomes/species/individual). We observed a highly significant correlation between *SR_j_* across the two datasets (p=3.30 × 10^−5^, Spearman correlation; Fig. 1B) providing an independent validation of the *SR_j_* estimates.

**Figure 1.**
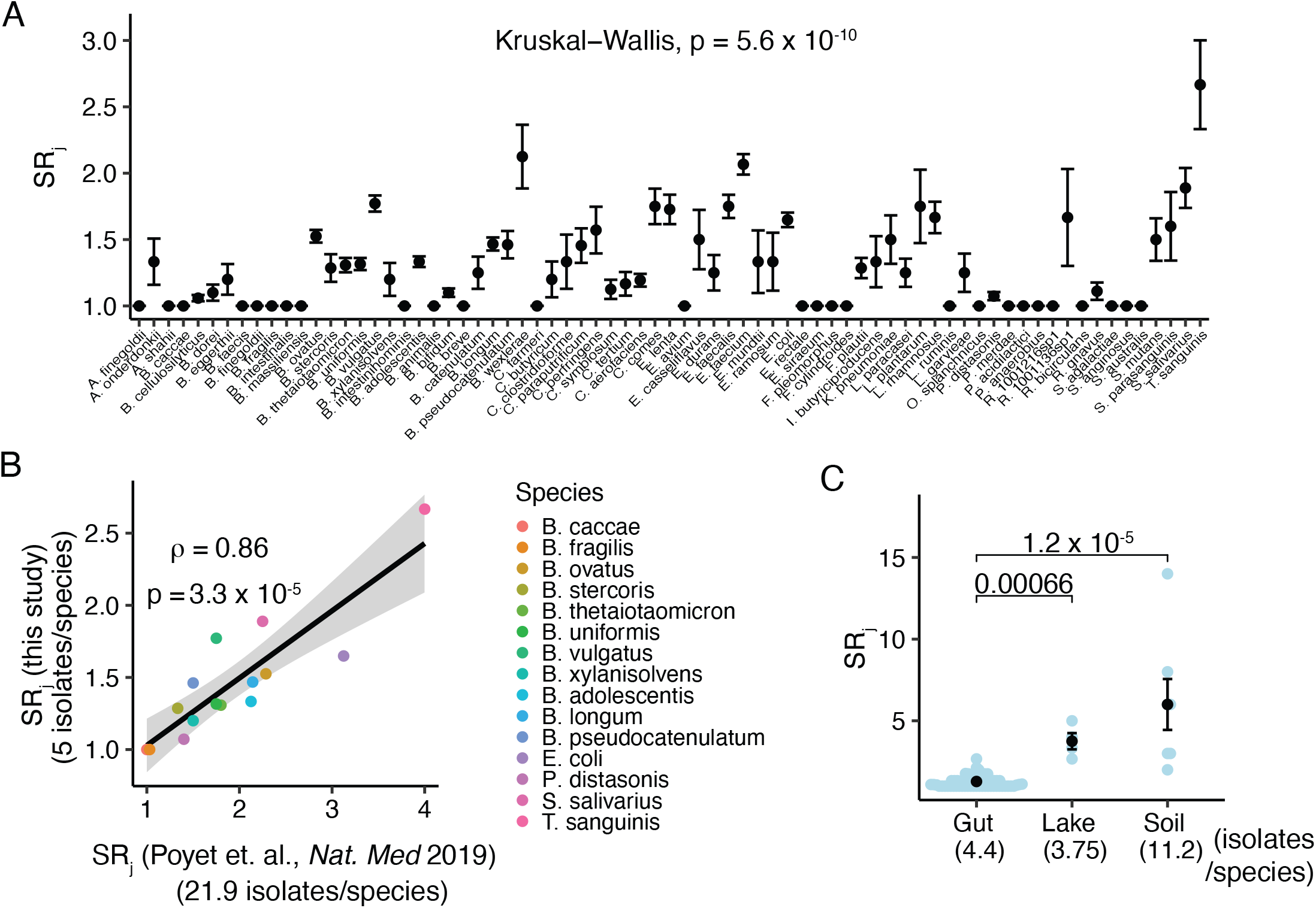
Strain richness (*SR_j_*) of human gut species. (A) *SR_j_* varies by species. (B) *SR_j_* using the isolate genome set in this study is highly correlated with *SR_j_* estimated from an independent set of bacterial genomes isoalted from 10 humans (Poyet et al., Nat Med 2019). (C) The *SR_j_* of human gut species is lower than *SR_j_* of species isolated from lake and soil microbiomes (Kruskal-Wallis test). Each blue point represents the average *SR_j_* for a species found in each of the environments. Black points represent the mean of the environment. Error bars indicate SEM.

Host colonization by multiple strains of a species within an individual’s microbiota could result from multiple independent strain colonization events or from diversification of one initial strain over a long time. We examined the distribution of pairwise distances between conspecific strains within versus between individuals and found that they varied widely and were not skewed towards more similar strains within an individual. Our results suggest multiple colonization events contributed to the presence of multiple strains within a person’s microbiota (Fig. S1G). When comparing the mean pairwise distances between species with *SR_j_* > 1 versus for species with *SR_j_* = 1, we found significantly greater pairwise distances in species with *SR_j_* > 1 compared to species with *SR_j_* = 1 (0.35 ± 0.001 versus 0.30 ± 0.005; p=2.2 × 10^−16^, Kruskal-Wallis), suggesting that larger accessory genomes in species with *SR_j_* > 1.00 might drive greater strain richness within individual microbiotas (Fig. S1H). In IBD, the gut microbiome has reduced genus and species diversity compared to healthy controls^28–31^. To determine if *SR_j_* is also altered in these states, we compared the average strain richness of species found in both healthy and disease microbiotas. We found that healthy and CD microbiomes do not differ in their strain counts at the species level whereas UC microbiomes may have lower *SR_j_* when compared to healthy microbiomes (Fig. S1I-J).

Finally, to determine how the strain richness structure of the human gut compares to that of other ecosystems, we compared the *SR_j_* of the human gut microbiome to that of soil and lake microbiomes. Species in the human gut demonstrated much lower strain richness (1.28 ± 0.33) than species sampled from the soil (6.00 ± 4.12) or lake (3.75 ± 1.00; Fig. 1C). This startling difference suggests that unique features of the gut, such as its semi-closed ecosystem with unidirectional flow, stable temperature, and fast microbial growth rates, may limit strain richness.

### Transmissibility and stability of strain richness

Previously, we demonstrated that the majority of donor microbiota strains stably engraft post-FMT in patients with recurrent *C. difficile* infection^8^, suggesting that donor strain richness might be transmitted via FMT. We isolated and sequenced 1,008 unique strains from 7 FMT donors and 13 recurrent *C. difficile* patients. We used the *Strainer* strain tracking algorithm to quantify the presence of donor strains in the recipients using the donor strain genomes and recipient metagenomes^8^. The *SR_j_* estimated from FMT donor metagenomics data closely resembled the *SR_j_* that we measured from cultured isolates in our complete cohort in Fig. 1 (Fig. S2A). This allowed us to track the transmission of donor strain-level community structure.

In this FMT study, single donors (Fig. 2A) were used for each recipient with six donors giving stool to six recurrent *C. difficile* recipients and one donor (D283) giving stool to seven different recurrent *C. difficile* recipients for a total of thirteen 1:1 donor-recipient pairs^32^. We quantified the number of donor strains per species found in the donors and the number of donor strains that were subsequently detected in recipients 8-weeks post-transplant (Fig. 2B, S2C). Donor *SR_j_* and the recipient *SR_j_* at week 8 post-FMT were significantly correlated (p = 4.20 x 10^-6^; Spearman’s rank correlation; Fig. 2C). Average *SR_j_* in recipients at week 8 post-FMT showed very little loss of donor strains and resembled the cultured *SR_j_* we initially measured in our cohort (Fig. S2B, S2D). Together these results demonstrate that a healthy donor strain-level community structure is transmissible via FMT.

**Figure 2.**
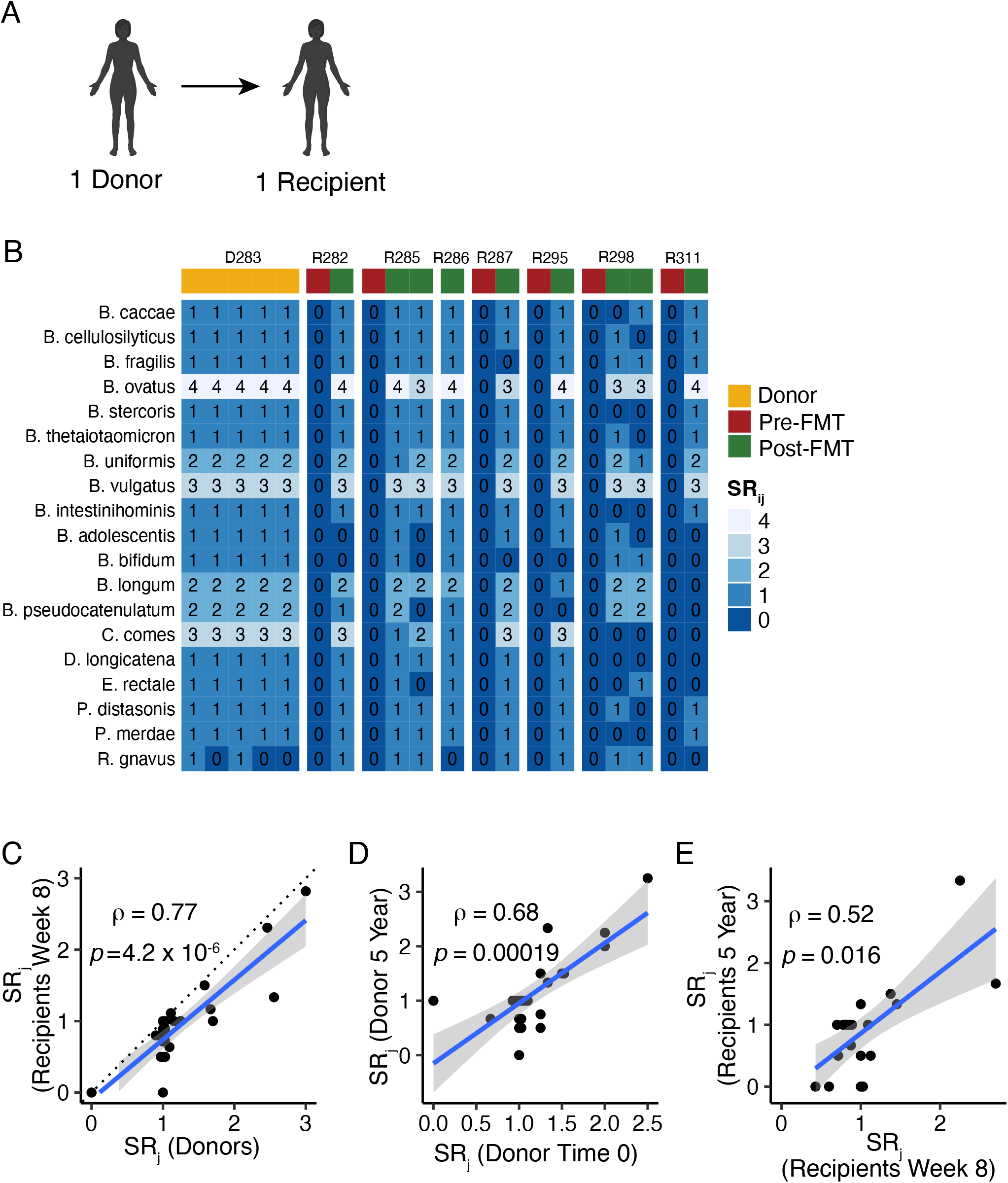
FMT durably transmits healthy donor *SR_j_* to recurrent *C. difficile* patients. (A) Schematic of FMT experimental design with 1:1 donor-recipient pairs. (B) Representative heatmap showing the transmission of *SR_ij_* from donor D283 to seven different recipients (R282, R285, etc…). The numbers in the cells of the heatmap indicate the number of strains of the given species detected in the donor or recipient across 5 time points for donor D283 and up to 3 time points for the 7 recipients. (C) Spearman correlation between recipient *SR_j_* at week 8 post-FMT and donor *SR_j_*. (D) Spearman correlation between donor *SR_j_* at 5 years and time 0 (pre-FMT stool sample). (E) Spearman correlation between recipient *SR_j_* at 5 years post-FMT and 8 weeks post-FMT. (C)-(E) Each point represents the average SR_j_ for the tracked species in (B).

A subset of donors (n=3) and recipients (n=3) from this FMT cohort had samples collected 5-years post-transplant which allowed us to measure the stability of strain richness in each species over time. In the donors, we find a significant correlation between donor *SR_j_* at timepoint 0 and 5 years later (Fig. 2D), suggesting long term stability of *SR_j_* in healthy, untransplanted individuals up to 5 years. In the recipients, we also find significant correlations between recipient *SR_j_* at 8 weeks post-FMT and 5-years post-FMT (Fig. 2E), suggesting that FMT can durably restore a healthy strain-level structure to recipients.

### Supraphysiologic strain richness administered to FMT recipients converges to the population average

Both the analyses of the unperturbed gut microbiota and of the transmission of the unperturbed gut microbiota suggest that strain richness ranges from approximately 1-3 strains per species and can be durably transmitted via FMT. However, we still do not know how elastic strain richness is, what the upper limits of *SR_j_* are, and what role ecologic and environmental pressures play in strain richness in the gut, as aspects such as modern antibiotic and hygiene practices may artificially reduce strain richness below any physiologic or ecologic limit. Pooled donor FMT offers a unique opportunity to test the ecologic limit of strain richness and determine if strain richness can be stably increased.

In the FOCUS clinical trial of FMT for UC patients, fourteen donor stools were combined into twenty-one different donor stool batches consisting of 3-7 unique donor stools per batch (Fig. 3A). Each multi-donor batch was given to one or more recipients 19 initially via colonoscopy and then at least 40 doses by enema^19^. For species that are common across healthy individuals, the pooled FMT approach provides a unique situation where a supraphysiologic number of strains for many species—much higher than what we found in our calculation of *SR_j_* in untransplanted individuals (Fig. 1A; Table 1)—was administered to the recipient. Thus, this is an excellent occasion to query the upper limits of *SR_j_* and its stability in recipients over time.

**Figure 3.**
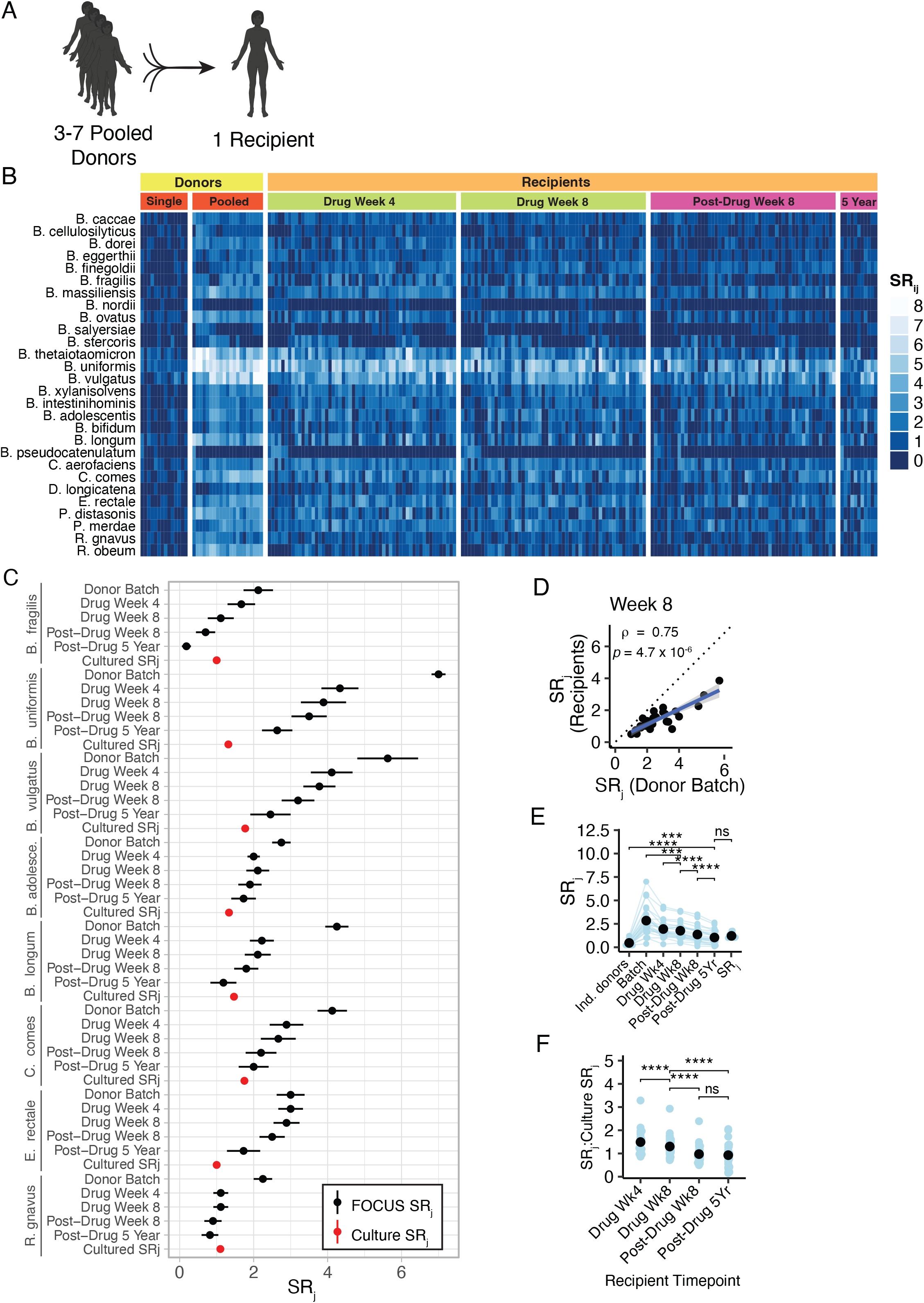
Supraphysiologic manipulation of strain richness in FMT recipients converges to the population baseline observed in untransplanted individuals. (A) Schematic for FMT experimental design with multi-donor stool batches administered to each recipient. (B) Heatmap showing the *SR_ij_* of single donors, donor batches, and recipients at post-FMT timepoints both during and after FMT drug administration. (C) *SR_j_* for representative species across the donor batch, recipient post-FMT timepoints, and cultured cohort from Fig. 1A (individual donors). (D) Spearman correlation between recipient *SR_j_* at drug week 8 and donor batch *SR_j_*. (E) *SR_j_* across donors, batches, recipient timepoints, and culture (previously measured in Fig. 1A and Table 1). (F) Ratio of Recipient *SR_j_*: Culture *SR_j_* across recipient post-FMT timepoints.

****p<10^-4^, Wilcoxon test, each blue point represents the ratio of recipient SR_j_:Culture SR_j_ of a tracked species. Black points represent the mean of the overall group. ns: not significant by Wilcoxon test.

Using *Strainer*, we quantified the number of donor strains present in the individual donors, in the donor batches, and in the recipient pre- and post-FMT timepoints. As with FMT subjects in the rCDI trial, we found that *Strainer* quantification of *SR_j_* across the individual donors (Fig. S3A) was significantly correlated with the *SR_j_* measured across our complete cohort in Fig. 1A, which included the FOCUS donors. As expected, the multi-donor pools harbored a higher *SR_j_* than in individuals (2.65 ± 1.27 vs. 1.28 ± 0.33; Fig. 3B-C).

After 8 weeks of transplantation, we find a strong correlation between the *SR_j_* of the donor batch and the *SR_j_* of the recipients (Fig. 3D). However, even after an intensive regimen of 40 fecal transplants over 8 weeks, only a proportion of donor *SR_j_* engrafted in recipients resulting in recipient *SR_j_* that was significantly less than the theoretical maximum based on the number of detected strains in the donor batches (1.33 ± 0.75 vs. 2.65 ± 1.27; p = 4.00 × 10^−6^, pairwise Wilcoxon signed-rank). These results demonstrate that even an intensive course of supraphysiologic strain richness via FMT for forty times over the course of eight weeks has a limited effect on the strain richness in the recipients.

When we track recipient *SR_j_* longitudinally, we observe a significant, progressive decrease in *SR_j_* in the recipients after the last FMT dose at week 8 (Fig. 3B-C). By 5 years after the final transplant, *SR_j_* in the recipients converged near the population wide *SR_j_* measured in individuals not receiving fecal transplants (Fig. 3C and Fig. 3E-F, Table S3). We found this trend to be consistent across all species, irrespective of species prevalence or *SR_j_* (Fig. S3B-G). Together, these results demonstrate that it is possible to therapeutically increase *SR_ij_* slightly and that there may be a limit near a strain richness of 5.00 (Fig. 3E). Additionally, this increase in recipient *SR_j_* is temporary as recipient *SR_j_* returns to the baseline *SR_j_* that we quantified in unperturbed healthy and disease microbiotas (Fig. 3E). These results suggest that strain richness in an individual has a species-specific capacity that is limited by the gut microbiota ecosystem.

## Discussion

Using high-throughput bacterial culturing and sequencing coupled with metagenomics and strain tracking, we find that the average strain richness of species in the healthy human gut microbiome varies by species and ranges between 1.00-2.67 (Fig. 1A). Previous investigations of microbiota composition in individuals with IBD at genus and species levels^28–30^ have shown reduced diversity compared to healthy individuals. We find that there is little to no difference in strain richness between healthy and CD microbiotas and a potentially lower strain richness in UC versus healthy (Fig. S1I-J). Comparing the strain richness of the human gut microbiome to environmental microbiomes demonstrated that this low level of bacterial strain diversity is unique to the semi-closed human gut ecosystem (Fig. 1C).

The factors that influence whether conspecific strains can co-exist within the gut microbiota are still being discovered. It is known that the multiple species of the gut microbiota mediate colonization resistance to enteric pathogens through interspecies inhibition of growth, direct killing, or production of bacteriocins^33–39^. Within a species, competition for nutrient resources becomes a major factor in colonization resistance as conspecific strains often have overlapping ecological niches^40^. For example, some less virulent *C. difficile* strains can decrease germination of enterotoxigenic *C. difficile* strains by competing for limited amino acid resources^41^. Another example is *B. fragilis*, which 16 expresses colonization factors that inhibit colonization by new conspecific strains^16^. Additionally, one recent study suggested that species that displayed strong codiversification with humans tended to have reduced genomes^42^, potentially resulting in a narrower range of niche utilization and increased competition. Species found to display strong codiversification with humans included *Collinsella aerofaciens* and *Eubacterium rectale* which we found to have a strain richness of 1.22 and 1.00, respectively. It is likely that these interspecies and intraspecies interactions along with other host-microbial interactions involving host immunity and codiversification apply selective pressures to certain bacterial species resulting in low richness at 1.0.

While most of the bacterial species we measured had low richness, there were also species that had strain richness ranging from 2.0-3.0 (Fig. 1A). Many open questions remain about the origins and mechanisms underlying the co-existence of multiple strains within a species. Our investigations into microbial comparative genomics offer some hints to underlying mechanisms. Comparing pairwise k-mer distances between conspecific strains, we observe similar distributions of pairwise distances within and across individuals (Fig. S1G). This distribution is consistent with the hypothesis that co-colonization of an individual with multiple strains within the same species arise from multiple colonization events and concur with the results of a recent study that found that strain replacements (i.e., separate colonization events over time) dominate the long-term (20-year timescale) landscape of genomic changes in commensals^43^. We also observe that conspecific strains from species with higher strain richness (*SR_j_* > 1.00 have significantly greater mean pairwise genomic distances compared to strains from species with low strain richness (*SR_j_* = 1.00), suggesting that larger accessory genomes confer a greater ability to utilize diverse niches and may facilitate greater strain richness within the human gut. One additional parameter that may play a role in facilitating higher strain richness is gut anatomy. One recent study on the human skin microbiome found that skin pores impose random bottlenecks as *Cutibacterium acnes* migrates into pores, reducing intraspecies competition and enabling coexistence of multiple *C. acnes* lineages^9^. In the gut, though the bulk of the niche volume is in the well-mixed lumen (allowing mixing and competition), anatomic features such as intestinal crypts may also promote co-colonization by multiple strains by partitioning a species population and reducing competition for shared resources.

Limitations to this study are that although the gut microbiota is relatively stable there is still a small amount of strain acquisition and loss that likely continually occurs in an individual. The duration of these transient colonizations will vary, but our pooled donor FMT results suggest that it can take weeks to years for therapeutically inflated strain richness of a species to converge. However, in untransplanted individuals, a transient increase in strain count is likely limited to a few species and different between individuals. Therefore, the impact of these dynamics on average strain richness for a species is likely minimal when averaged across more than 60 individuals. Another limitation is that there is no current tool even close to being able to evaluate every bacterial cell in a person’s gut microbiome at the strain level. Therefore, we cannot know if there is a large amount of strain diversity at very low abundance that is undetectable with current methods. Although our deeper sampling of a few species suggested we had not found all strains of a species in every person, the overall impact of deeper sampling was minimal. In addition, our estimates of strain richness were confirmed by an independent dataset^27^. Finally, there is not a widely-accepted definition of a bacterial strain. However, we used a genomics-based threshold that was empirically demonstrated to identify bacterial strains shared over time^20^, in between family members^20^, and across fecal transplants^8^. Our analyses in the set of bacterial genomes in this analysis further support a strain threshold of around 0.96 kmer overlap (Fig. S1A).

Despite many potential facilitators for increased strain diversity, our results demonstrate that the human gut ecosystem does not permit limitless strain richness. When pooled donor FMT is used to administer supraphysiologic strain richness to recipients, we find only a temporary increase in recipient strain richness that eventually returns to the population baseline (Fig. 3B-C, Fig. 3E-F). In one-to-one donor-recipient pairs, however, donor strain richness is consistently transmittable and stable within recurrent *C. difficile* recipients (Fig 2B-C). Together, these findings emphasize the importance of understanding the influence of human gut anatomy and physiology on strain richness and its immediate translational applications. In FMT and defined live biotherapeutic products, where success depends on the successful engraftment of bacterial strains, careful consideration of which strains to include, how many strains of a species to include, and whether resident strains should be removed is essential to future development. A recent meta-analysis found recipient resident species to play an outsize role in inhibiting donor strain engraftment^42^. Together with the limited strain-richness capacity of the gut, these data provide a clear theoretical and practical basis for removing risk- or disease-associated strains from the gut by administration of other milder strains from the same species that are more fit and occupy the same species niche. Likewise, these results also suggest that attempts to dose supraphysiologic strain richness to recipients would require continuous administration or should expect only a temporary increase with a lower strain richness remaining in the long term.

## Methods

### Human participants

For the recurrent CDI study, written consent was obtained from all individuals recruited in the study using a protocol approved by the Mount Sinai Institutional Review Board (HS no. 11-01669). Donors and patients who received FMT for recurrent CDI or recurrent CDI and IBD were described in a previous study analyzed with 16S ribosomal RNA amplicon sequencing^32^. For the Fecal Microbiota Transplantation for Chronic Active Ulcerative Colitis (FOCUS) study, written informed consent was obtained from all patients prior to screening. Donors and patients were who received FMT for UC were described in a previous study^19^. Additional patients were recruited at The Mount Sinai Hospital under IRB 16-00021. For analysis, we only considered for analysis the subset of individuals for which donor and recipient stool samples from multiple time points had been collected.

### Fecal sample collection and high-throughput anaerobic bacterial isolation

We followed the protocol previously described in^4,44,45^. Briefly, fecal samples were aliquoted on dry ice or liquid nitrogen and stored at −80°C. Under strict anaerobic conditions, stool from each donor was blended into culture media^44^ and stored at −80°C. We utilized a well-established, robotized platform that enables isolation and culturing of a high proportion of bacteria found in the human gut^3,4,6,20,44^. Briefly, clarified and diluted stool donor stool was plated onto a variety of solid selective and non-selective media under anaerobic, micro-aerophilic, and aerobic conditions selected to promote the greatest growth of a diverse array of all stool microbes. Plates were incubated for 48-72 hours at 37°C. 384 single colonies from each donor microbiota were individually picked and regrown in media for 48 hours under anaerobic conditions. Regrown isolates were identified at the species level using a combination of MALDI-TOF mass spectrometry (Bruker Biotyper) and 16S rDNA amplicon sequencing. The original 384 isolates were then de-replicated and approximately three isolates of each species were archived in multi-well plates. DNA was then extracted by bead-beating and extraction in phenol chloroform and stored at −20°C. Across these culture collections, we have found an individual’s cultured strains represent the majority of bacteria assigned reads in the metagenome with approximately 70% of the bacterial metagenome mapping to the cultured strain genomes^8^.

### Strain enrichment in gnotobiotic mice

To potentially enrich for strains at lower abundance in the human donor, we colonized ex-germ free mice with the same human stool samples previously used for direct culturing and administered two different diets to the colonized mice. We selected the two diets (41% high casein, Harlan TD.09054 and 5% psyllium, Harlan TD.150229) from our published screen of >40 custom mouse diets^46^. We collected fecal pellets after two weeks of diet administration and stored the them at −80°C. Mouse fecal pellets were then used for depth-focused high-throughput culture to more deeply sample the strain richness of our selected species (Table S1).

### Selection of species for SR_j_ validation and depth-focused high-throughput anaerobic bacterial culture

We selected ten gut bacterial species for their membership across the four major phyla of the gut and for their range of *SR_j_* based on preliminary calculations. To enable greater sampling depth for these species, we adjusted our previous breadth-focused culturing approach by plating clarified stool samples on a selected range of environmental conditions designed to cultivate our target species. Next, 384 colonies were picked for each donor sample and regrown in liquid media in multi-well plates. Each isolate was then identified by a combination of matrix-assisted laser desorption/ionization-time of flight mass spectrometry and whole-genome sequencing. Using this knowledge, the original 384 isolates were de-replicated and approximately ten isolates of each of the target species were archived in multi-well plates. DNA was then extracted by bead beating and stored at −20°C.

### Construction of whole genome libraries and Illumina sequencing

Illumina library construction for performed using the seqWell Plexwell 384 kit. DNA was barcoded, ligation products were purified, and finally an enrichment PCR was performed. Samples were pooled in equal proportions and size-selected before sequencing with an Illumina HiSeq (paired-end 150 base pairs (bp)). The sequence data files (FASTQ) for all whole genome assemblies are available on NCBI (BioProject ID: PRJNA880610).

### Strains as bacterial isolates with <96% similarity, metagenomic strain tracking, and calculation of SR_j_

As in our previous analyses^3,4,20^, bacterial isolates with less than 96% whole-genome similarity were defined as unique strains, otherwise they were considered as multiple isolates for the representative strain. To calculate an average strain richness for a given species from cultured isolates, *SR_j_*, we calculated the strain richness of a species *j* within an individual *i*: *SR_ij_*. We then average *SR_ij_* across the individuals in our cohort to arrive at an average *SR_j_*. We only measure *SR_ij_* where an individual harbored species *j* within their microbiota. Thus, *SR_j_* is a measure of the number of strains a species stably maintains if it is present within a microbiota. For our metagenomics analyses, we used our previously published metagenomics algorithm *Strainer^8^*; we tracked the presence or absence of a strain in a sample for quantification of *SR_j_*.

### Strainer algorithm for detection of cultured and sequenced strains from metagenomics samples

We previously described *Strainer*^8^ for tracking discrete bacterial strain genomes in metagenomes. In brief, we identify a set of informative sequence features, or k-mers, from a bacterial genome that can uniquely identify any given strain. We first initialize this informative k-mer set by removing those shared extensively with bacterial genomes and fecal metagenomes from unrelated, non-cohabitating individuals, where the probability of the occurrence of the same strain is very low. Next, we update this informative k-mer set by removing those that co-occur on metagenomics reads with uninformative k-mers. Finally, we assign each sequencing read in a metagenomics sample of interest to a unique strain by comparing the distribution of *k*-mers on a read with the informative *k*-mers identified earlier, and with controls to find statistical significance. In the recurrent *C. difficile* trial, we applied *Strainer* on metagenomics samples from 7 donors and 13 recipients over multiple timepoints (pre-FMT to 5-years post-FMT). We sequenced an average of approximately 5.2 million reads from a total of 85 metagenomics samples across donors and recipients. For the FOCUS UC trial, we also applied *Strainer* on metagenomics samples from 14 donors, 21 pooled batch samples and 63 recipients over multiple timepoints (pre-FMT to 5 years post-FMT). The average sequencing depth of metagenomics was 2.4 million reads and we tracked 1421 unique strains from the donors in the recipients. In the FOCUS UC trial, treated subjects were either given an endoscopic FMT followed by 40 enemas over 8 weeks (active arm of placebo-controlled trial) or 40 enemas over 8 weeks without endoscopically administered FMT (optional open-label arm). Both groups were combined for our analyses. For understanding the impact of pooled donors on strain-richness limits, we focused on strains that engrafted in at least 30% of recipients.

### Statistical analysis and plotting

Analysis was performed in RStudio v2022.02.3+492. Heatmaps were created using the R packages ComplexHeatmap v2.12.0 and circlize v0.4.15.

## Acknowledgements

This work was supported in part by the staff and resources of the Mount Sinai Gnotobiotic Facility and the Scientific Computing Division at the Icahn School of Medicine at Mount Sinai. We thank C. Fermin, E. Vazquez, and G.N. Escano for gnotobiotic husbandry. This work was supported by the National Institutes of Health grants (nos. NIDDK DK112978, NIDDK DK124133, NIDDK DK123749), an NIH F30 to A.C. (NIDDK DK131862), a Crohn’s and Colitis Foundation RFA award to V.A. (no. 650451), and Janssen Research & Development.

## Figure Legends

**Figure S1.**
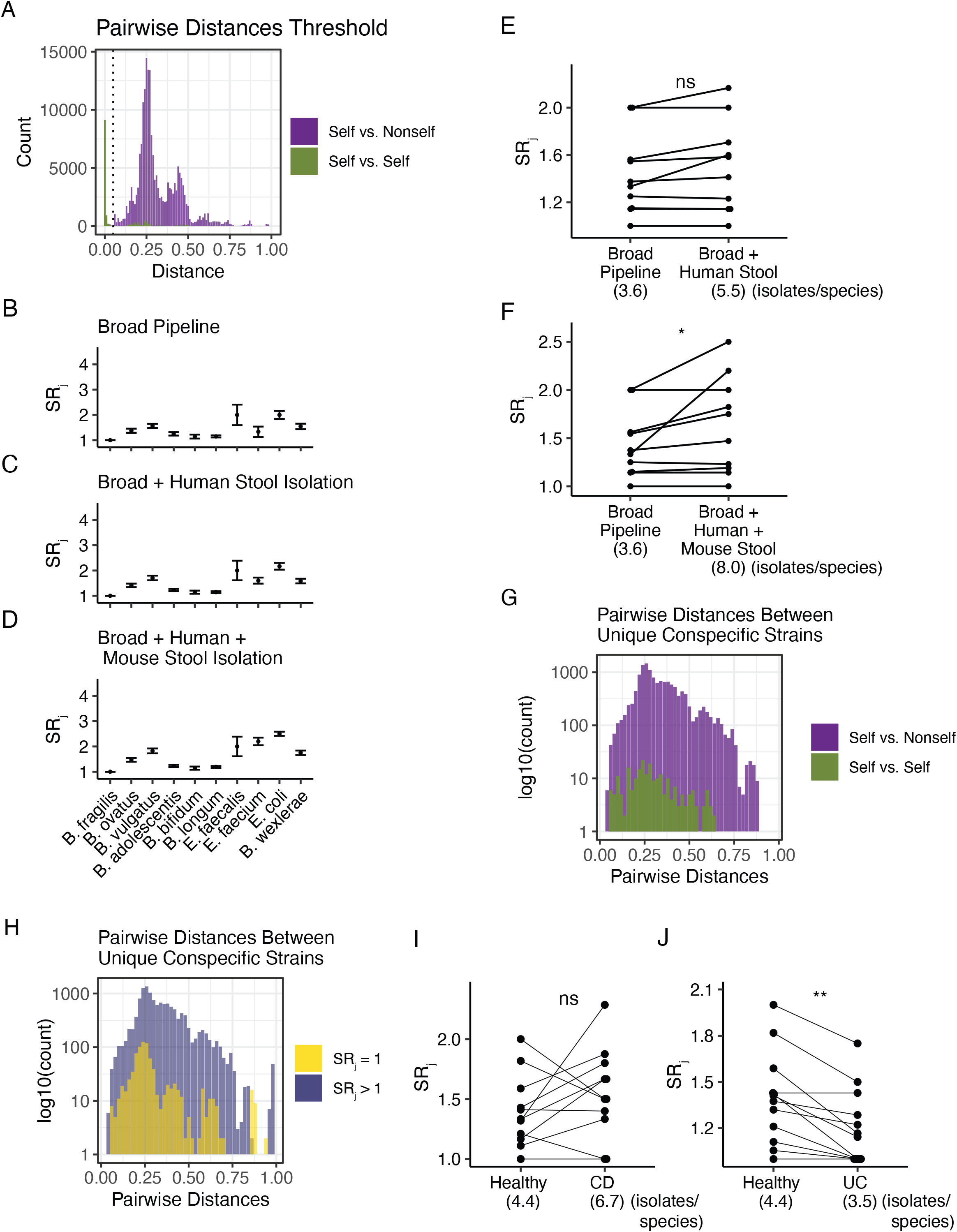
Determination of *SR_j_* strain threshold and validation of *SR_j_* with deeper sampling of a subset of cultured gut bacterial species. (A) Differences in kmer content were calculated for all pairwise combinations of isolates from a species cultured from two unrelated individuals (purple) with no direct microbial transfer between them or an individual’s own microbes from a single timepoint (green). Dotted line shows the threshold of 0.05 kmer difference. (B) Preliminary calculation of *SR_j_* using genomes isolated from the Broad pipeline (standard pipeline used to create libraries of cultured gut bacteria). For deeper sampling estimates of *SR_j_*, we isolated additional genomes from the same species from (C) the original human stool sample and (D) mouse stool samples from gnotobiotic mice colonized with the same human stool and given unique diets for strain enrichment. Comparison of *SR_j_* calculated with genomes from the broad pipeline or in combination with (E) human stool samples, or (F) human and mouse stool samples. (G) Distribution of pairwise kmer distances between unique conspecific strains across and within individuals. (H) Distribution of pairwise kmer distances between unique conspecific strains of species across low and high *SR_j_*. Paired species comparisons of healthy *SR_j_* versus (I) Crohn’s Disease and (J) ulcerative colitis. *p<0.05, paired Wilcoxon test, each point represents the average *SR_j_* of one of the target species.

**p<0.01, paired Wilcoxon test, each point represents the average *SR_j_* measured from microbiomes of a healthy or disease state.

ns: not significant by paired Wilcoxon test.

**Figure S2.**
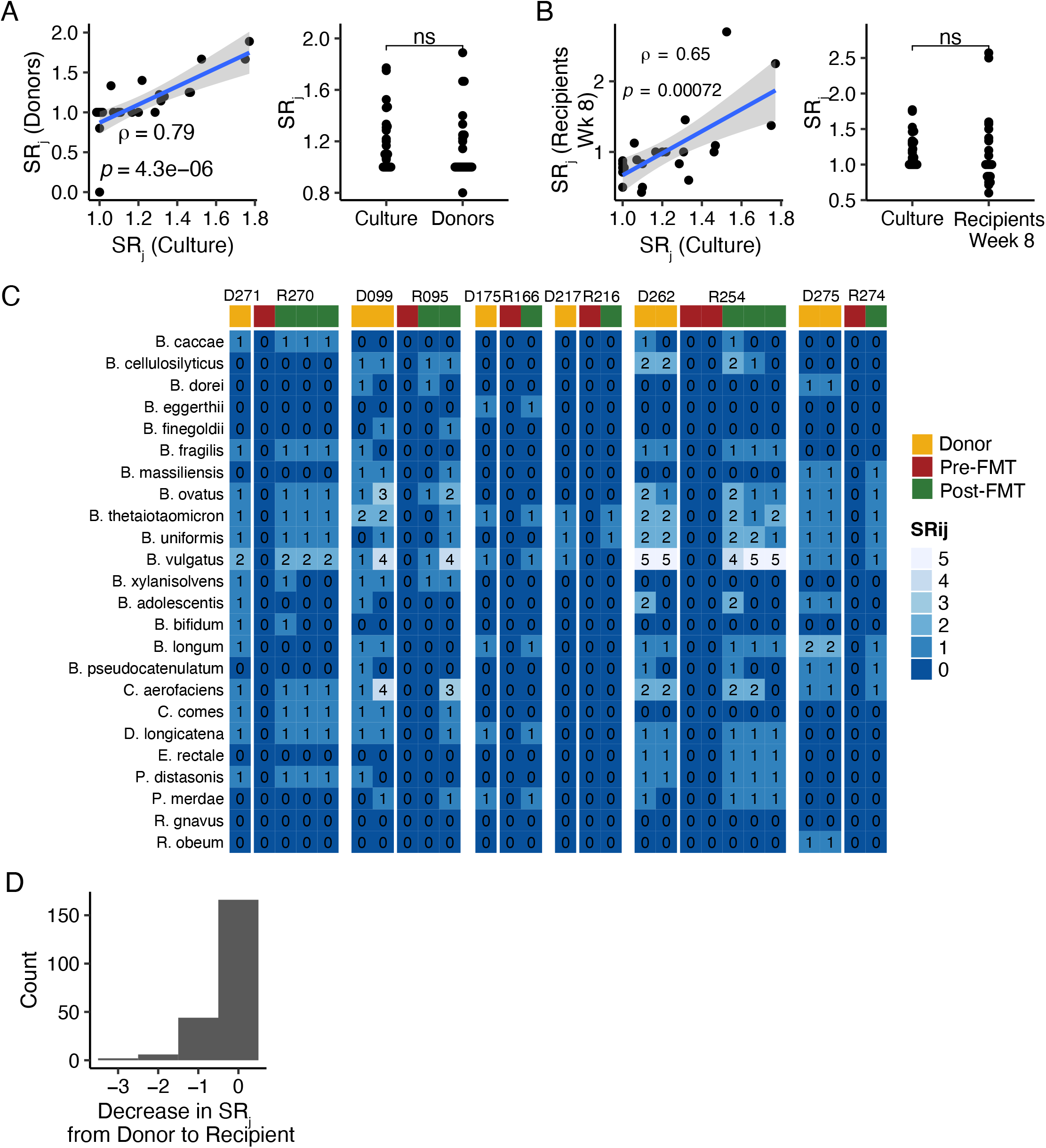
Metagenomics-quantified *SR_j_* correlates with the cultured *SR_j_* measured across our cohort. (A) Spearman correlation between donor *SR_j_* as measured by metagenomics and *SR_j_* from our original cultured cohort (first panel) and differences in *SR_j_* between the two groups (second panel). (B) Spearman correlation between recipient *SR_j_* as measured by metagenomics at week 8 post-FMT and *SR_j_* from our original cohort (first panel) and differences in *SR_j_* between the two groups (second panel). (A) and (B) Each point represents the average *SR_j_* for species. (C) Heatmap representing the remaining six donors who donated stool to six different recipients. (D) Distribution of change in strain richness from donor to recipients at week 8 post-FMT.

**Figure S3.**
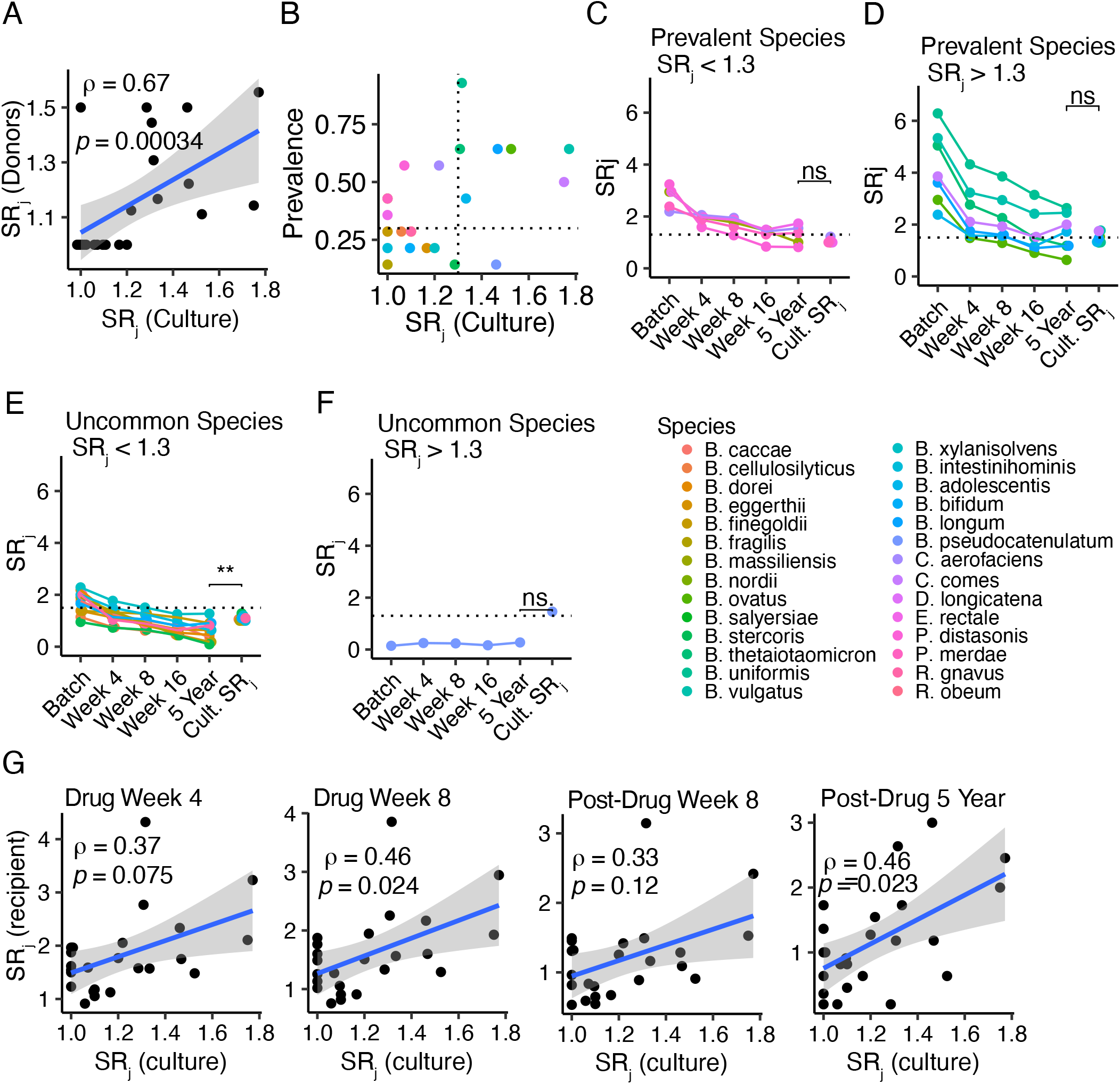
SR_j_ trends towards baseline over time. (A) Spearman correlation between the metagenomics SR_j_ of the individual FOCUS donors with the overall cultured SR_j_ measured across our cohort. (B) Bacterial species prevalence vs cultured *SR_j_*. Dotted lines show cutoffs for determining high versus low prevalence (30%) and high versus low *SR_j_* (1.30). Prevalence for each species was calculated based on the number of donors that harbored that species divided by the total number of donors. Average *SR_j_* at each timepoint for (C) prevalent species with *SR_j_* < 1.3, (D) prevalent species with *SR_j_* > 1.3, (E) uncommon species with *SR_j_* < 1.3, and (F) uncommon species with *SR_j_* > 1.3. (G) Spearman correlation between recipient *SR_j_* and culture *SR_j_* across recipient post-FMT timepoints.

**p<0.01, Wilcoxon test.

ns: not significant by Wilcoxon test

## References

1. Parida, S. et al. A Procarcinogenic Colon Microbe Promotes Breast Tumorigenesis and Metastatic Progression and Concomitantly Activates Notch and ß-Catenin Axes. Cancer Discov 11, 1138–1157 (2021).

2. Arthur, J. C. et al. Intestinal inflammation targets cancer-inducing activity of the microbiota. Science 338, 120–123 (2012).

3. Britton, G. J. et al. Defined microbiota transplant restores Th17/RORγt+ regulatory T cell balance in mice colonized with inflammatory bowel disease microbiotas. Proceedings of the National Academy of Sciences of the United States of America 117, 21536–21545 (2020).

4. Yang, C. et al. Fecal IgA Levels Are Determined by Strain-Level Differences in Bacteroides ovatus and Are Modifiable by Gut Microbiota Manipulation. Cell Host & Microbe 27, 1–9 (2020).

5. Yang, C. et al. Immunoglobulin A antibody composition is sculpted to bind the self gut microbiome. Science Immunology 7, eabg3208 (2022).

6. Spindler, M. P. et al. Human gut microbiota stimulate defined innate immune responses that vary from phylum to strain. Cell Host & Microbe 0, (2022).

7. Zhao, S. et al. Adaptive Evolution within Gut Microbiomes of Healthy People. Cell Host and Microbe 25, 656–667.e8 (2019).

8. Aggarwala, V. et al. Precise quantification of bacterial strains after fecal microbiota transplantation delineates long-term engraftment and explains outcomes. Nature Microbiology 6, 1309–1318 (2021).

9. Conwill, A. et al. Anatomy promotes neutral coexistence of strains in the human skin microbiome. Cell Host & Microbe 30, 171–182.e7 (2022).

10. Olm, M. R. et al. inStrain profiles population microdiversity from metagenomic data and sensitively detects shared microbial strains. Nat Biotechnol 39, 727–736 (2021).

11. Drewes, J. L. et al. Transmission and clearance of potential procarcinogenic bacteria during fecal microbiota transplantation for recurrent Clostridioides difficile. JCI Insight 4, 130848 (2019).

12. Vatanen, T. et al. Genomic variation and strain-specific functional adaptation in the human gut microbiome during early life. Nature Microbiology 4, 470–479 (2019).

13. Zheng, W. et al. High-throughput, single-microbe genomics with strain resolution, applied to a human gut microbiome. Science 376, eabm1483 (2022).

14. Gupta, S. et al. LB973 Cutaneous surgical wounds have distinct microbiomes from intact skin. Journal of Investigative Dermatology 142, B24 (2022).

15. Blaser, Martin. Missing Microbes: How the Overuse of Antibiotics is Fueling Our Modern Plagues. (2014).

16. Lee, S. M. et al. Bacterial colonization factors control specificity and stability of the gut microbiota. Nature 501, 426–431 (2013).

17. Verster, A. J. et al. The Landscape of Type VI Secretion across Human Gut Microbiomes Reveals Its Role in Community Composition. Cell Host & Microbe 22, 411–419.e4 (2017).

18. Yassour, M. et al. Natural history of the infant gut microbiome and impact of antibiotic treatment on bacterial strain diversity and stability. Science Translational Medicine 8, 343ra81 (2016).

19. Paramsothy, S. et al. Multidonor intensive faecal microbiota transplantation for active ulcerative colitis: a randomised placebo-controlled trial. The Lancet 389, 1218–1228 (2017).

20. Faith, J. J. et al. The long-term stability of the human gut microbiota. Science 341, 1237439 (2013).

21. Faith, J. J., Colombel, J.-F. & Gordon, J. I. Identifying strains that contribute to complex diseases through the study of microbial inheritance. PNAS 112, 633–640 (2015).

22. Faith, J. J. et al. Strain population structure varies widely across bacterial species and predicts strain colonization in unrelated individuals. bioRxiv 2020.10.17.343640 (2020) doi:10.1101/2020.10.17.343640.

23. Faith, J. J., McNulty, N. P., Rey, F. E. & Gordon, J. I. Predicting a human gut microbiota’s response to diet in gnotobiotic mice. Science 333, 101–104 (2011).

24. Bahtiyar Yilmaz, A. et al. Long-term evolution and short-term adaptation of microbiota strains and sub-strains in mice A B Yilmaz et al ll Resource Long-term evolution and short-term adaptation of microbiota strains and sub-strains in mice. Cell Host & Microbe 29, 1–14 (2021).

25. McNulty, N. P. et al. Effects of diet on resource utilization by a model human gut microbiota containing Bacteroides cellulosilyticus WH2, a symbiont with an extensive glycobiome. PLoS Biol 11, e1001637 (2013).

26. Walker, A. W. et al. Dominant and diet-responsive groups of bacteria within the human colonic microbiota. ISME J 5, 220–230 (2011).

27. Poyet, M. et al. A library of human gut bacterial isolates paired with longitudinal multiomics data enables mechanistic microbiome research. Nature Medicine 25, 1442–1452 (2019).

28. Forbes, J. D. et al. A comparative study of the gut microbiota in immune-mediated inflammatory diseases - Does a common dysbiosis exist? Microbiome 6, (2018).

29. Pascal, V. et al. A microbial signature for Crohn’s disease. Gut 66, 813–822 (2017).

30. Michail, S. et al. Alterations in the gut microbiome of children with severe ulcerative colitis. Inflammatory Bowel Diseases 18, 1799–1808 (2012).

31. Antharam, V. C. et al. Intestinal dysbiosis and depletion of butyrogenic bacteria in Clostridium difficile infection and nosocomial diarrhea. Journal of Clinical Microbiology 51, 2884–2892 (2013).

32. Hirten, R. P. et al. Microbial Engraftment and Efficacy of Fecal Microbiota Transplant for Clostridium Difficile in Patients With and Without Inflammatory Bowel Disease. Inflamm Bowel Dis 25, 969–979 (2019).

33. Sassone-Corsi, M. et al. Microcins mediate competition among Enterobacteriaceae in the inflamed gut. Nature 540, 280–283 (2016).

34. Becattini, S. et al. Commensal microbes provide first line defense against Listeria monocytogenes infection. Journal of Experimental Medicine 214, 1973–1989 (2017).

35. Caballero, S. et al. Cooperating commensals restore colonization resistance to vancomycin-resistant Enterococcus faecium. Cell Host Microbe 21, 592–602.e4 (2017).

36. Ducarmon, Q. R. et al. Gut Microbiota and Colonization Resistance against Bacterial Enteric Infection. Microbiol Mol Biol Rev 83, e00007–19 (2019).

37. Donia, M. S. et al. A systematic analysis of biosynthetic gene clusters in the human microbiome reveals a common family of antibiotics. Cell 158, 1402–1414 (2014).

38. Chatzidaki-Livanis, M., Coyne, M. J. & Comstock, L. E. An antimicrobial protein of the gut symbiont Bacteroides fragilis with a MACPF domain of host immune proteins. Mol Microbiol 94, 1361–1374 (2014).

39. Roelofs, K. G., Coyne, M. J., Gentyala, R. R., Chatzidaki-Livanis, M. & Comstock, L. E. Bacteroidales Secreted Antimicrobial Proteins Target Surface Molecules Necessary for Gut Colonization and Mediate Competition In Vivo. mBio 7, e01055–16 (2016).

40. Cohan, F. M. Transmission in the Origins of Bacterial Diversity, From Ecotypes to Phyla. Microbiol Spectr 5, (2017).

41. Leslie, J. L. et al. Protection from Lethal Clostridioides difficile Infection via Intraspecies Competition for Cogerminant. mBio 12, e00522–21 (2021).

42. Schmidt, T. S. B. et al. Drivers and determinants of strain dynamics following fecal microbiota transplantation. Nat Med 28, 1902–1912 (2022).

43. Garud, N. R., Good, B. H., Hallatschek, O. & Pollard, K. S. Evolutionary dynamics of bacteria in the gut microbiome within and across hosts. PLoS Biol 17, e3000102 (2019).

44. Britton, G. J. et al. Microbiotas from Humans with Inflammatory Bowel Disease Alter the Balance of Gut Th17 and RORγt+ Regulatory T Cells and Exacerbate Colitis in Mice. Immunity 50, 212–224 (2019).

45. Contijoch, E. J. et al. Gut microbiota density influences host physiology and is shaped by host and microbial factors. eLife 8, (2019).

46. Llewellyn, S. R. et al. Interactions Between Diet and the Intestinal Microbiota Alter Intestinal Permeability and Colitis Severity in Mice. Gastroenterology 154, 1037–1046.e2 (2018).

